# Cytoplasmic Aggregation of RPB1 Predicts Failure of Neoadjuvant Chemotherapy

**DOI:** 10.1101/2023.01.13.523908

**Authors:** Bence Nagy-Mikó, Orsolya Németh-Szatmári, Réka Faragó-Mészáros, Aliz Csókási, Bence Bognár, Nóra Ördög, Barbara N. Borsos, Hajnalka Majoros, Zsuzsanna Ujfaludi, Orsolya Oláh-Németh, Aliz Nikolényi, Ágnes Dobi, Renáta Kószó, Dóra Sántha, György Lázár, Zsolt Simonka, Attila Paszt, Katalin Ormándi, Tibor Pankotai, Imre M. Boros, Zoltán Villányi, András Vörös

**Author notes:** Correspondence: Zoltán Villányi, András Vörös. Declaration of competing interest: None.

## Abstract

We aimed to investigate the contribution of co-translational protein aggregation to the chemotherapy resistance of tumor cells. Increased co-translational protein aggregation reflects altered translation regulation that may have the potential to buffer transcription under genotoxic stress. As an indicator for such event, we followed cytoplasmic aggregation of RPB1, the aggregation prone largest subunit of RNA polymerase II, in biopsy samples taken from patients with invasive carcinoma of no special type. RPB1 frequently aggregates co-translationally in the absence of proper HSP90 chaperone function or in ribosome mutant cells as revealed formerly in yeast. We found that cytoplasmic foci of RPB1 occur in larger sizes in tumors that showed no regression after therapy. Based on these results, we propose that monitoring the cytoplasmic aggregation of RPB1 may be suitable for determining – from biopsy samples taken before treatment – the effectiveness of neoadjuvant chemotherapy.

## Introduction

Neoadjuvant chemotherapy is a common approach for treating breast cancer [1]. The predictive markers used today (Estrogen Receptor, Progesterone Receptor, Human Epidermal Growth Factor Receptor 2 and the Marker of Proliferation Ki-67) are very useful for selecting the appropriate drugs in each case; however, there is a lack of predictive markers for the expected outcome of the selected therapy. Epirubicin, an anthracycline topoisomerase II inhibitor that acts as a DNA intercalating agent thereby inhibiting both transcription and replication processes, is a widely administered drug in neoadjuvant chemotherapy [2]. Based on the effectiveness of the drug, tumors can be classified into the following three main groups: complete, partial, and non-regressive respectively. One possible explanation for epirubicin resistance is that cells rely on existing mRNA depositories under genotoxic stress when transcription is blocked. Stress granules and P-bodies are phase separated membraneless organelles that together with other RNA-protein complexes play important roles in the stress response of cells and maintain a transcription independent source of mRNAs [3]. The recently discovered NOT1-containing assemblysomes are another type of these RNP granules with a potential role in response of tumor cells to therapy [4,5].

Assemblysomes may play a role in stress response by supporting the co-translational assembly of stress-responsive protein complexes. One already confirmed complex dependent on assemblysomes for its integrity is the 26S proteasome, but according to recent *in silico* predictions complexes involved in DNA damage response are also regulated by assemblysomes [4,5]. Assemblysomes contain the NOT1 subunit of the CCR4–NOT complex, the major deadenylase of eukaryotic cells [4]. NOT1 has been recently reported to function as a chaperone platform (for reviews see [6–8]). Impairment of the chaperone capacity of the cell leads to failure in the assembly of protein complexes. Consequently, the loss of function of complexes playing role in apoptosis or cellular immune response processes can be beneficial for tumor progression. RPB1 is the largest subunit of the eukaryotic RNA polymerase II (RNAPII) and — as revealed in yeast — is a highly aggregation-prone protein that depends both on the CCR4—NOT complex and the chaperone HSP90 for its native conformation [9]. RNAPII is a 12 subunit complex. It assembles in the cytoplasm from RPB1 and RPB2 sub-complexes [10]. We hypothesized in a former study that if the CCR4-NOT complex plays a role in RPB1 and RPB2 sub-complex assembly than generating a condition where both a CCR4-NOT subunit and RPB2 are limiting, will lead to RPB1 accumulation in the cytoplasm as only the fully assembled RNAPII can bind to Iwr1 that facilitates nuclear import of the assembled complex [11]. According to our former results ovaries of NOT3 and RPB2 trans-heterozygous *Drosophila melanogaster* accumulated RPB1 foci in the cytoplasm revealing not only the role of the CCR4-NOT complex in RNAPII assembly but also that Rpb1 aggregation during CCR4-NOT mediated co-translational RNAPII assembly is a conserved phenomenon [9].

Tumors belonging to the non-regressive group in their response to neoadjuvant chemotherapy are resistant toward the toxic transcription inhibitory effect of epirubicin. We considered the possibility that the reason behind the phenomenon is that epirubicin resistant cells are able to compensate for the loss of transcription by buffering their gene expression relying on mRNA deposited in condensates such as P-bodies, stress granules or NOT1 containing assemblysomes. Since it has been recently reported that these cytoplasmic bodies co-localize with, or in the case of assemblysomes their formation is even dependent on NOT1 [4,12], it is reasonable to assume that increased P-body, stress granule, and assemblysome biogenesis may generate a condition where soluble NOT1 becomes limiting. This may lead at least partially to the loss of function of NOT1 in processes, including its chaperone platform role in the correct folding of RPB1 [9]. Considering that RPB1 is a component of RNAPII, the major machinery of transcription, it may be even permitted to lose significant active RPB1 in the form of aggregates in tumor cell clones that otherwise tolerate complete blockade of transcription by epirubicin. Therefore, Rpb1 cytoplasmic aggregation might be a potent indicator of tumor cell clones in biopsies that will tolerate transcription blockers used as chemotherapeutic agents.

## Materials and methods

### Cohort selection

A total of 25 patients diagnosed with invasive carcinoma of NST and 30 patients diagnosed with clear cell renal cell carcinoma, who underwent radical or partial nephrectomy at the University of Szeged, were selected for this study. This study was conducted with the permission of the Regional and Institutional Human Medical Biological Research Ethics Committee, University of Szeged (No. 188/2019-SZTE and 5181/2022-SZTE). It was conducted in accordance with the Declaration of Helsinki and approved by the Institutional Review Board (or Ethics Committee) of Science and Research Ethics of the Medical Research Council (protocol code: IV/5376-2/2020/EKU).

### Preparations of normal and tumorous tissues

For immunofluorescence staining, the tissues were embedded in Shandon Cryomatrix gel (Thermo Fisher Scientific), and 5 μm sections were prepared on Superfrost Ultra Plus slides by cryostat (Cryostar NX50).

### Preparations of formalin-fixed paraffin-embedded (FFPE) biopsy samples

Tissues were fixed with 10% formalin for 24—48 h at room temperature. The process of paraffin embedding used ethanol + IPA (70%-96%) and xylene/IPA, following standard protocols. Biopsy samples were embedded in paraffin blocks, and 5—μm sections were prepared using microtome following the protocol described in [13].

### Immunostaining of frozen tissues

Tissues were fixed for 10 min using acetone and subsequently washed three times with PBS solution. Subsequently, the tissues were permeabilized with 0.3% Triton-X-100/PBS for 20 min at 25°C. The sections were subsequently blocked with 5 % BSA/0.3 % Triton X-100/PBS for 1 h. The samples were incubated with primary antibodies diluted in 1 % BSA/PBST: anti-RNAPII (1BP7G5 from L. Tora, IGBMC) in 1:250 dilution. After the washing steps, secondary GAM Alexa 488 (Molecular Probes, A11029) antibody in 1:500 dilution was used. Finally, the cells were mounted with DAPI-containing ProLong Gold Antifade Reagent (Life Technologies). The samples were visualised using FLUOVIEW FV10i confocal microscopy. The same exposition time was used for every image capturing.

### Immunostaining of FFPE samples

Paraffin was removed by soaking slides for 10 min in xylene. Rehydrating was performed by soaking slides in sequentially decreasing v/v% ethanol (100%, 95%, 70% and 50%) for 2 min each. Samples were permeabilized for 15 min in 0.1% TritonX-100 at room temperature. Three washing steps were performed with PBS, 30 s each. Blocking was performed in 5% BSA/PBS for 1 h. Primary anti-RNAPII (Abcam, ab26721 or Santa-Cruz, sc-47701) was used in 1:200 dilution in 5% BSA/PBS. After the washing steps, secondary GAM Alexa 647 (Abcam, ab150115) or GAR Alexa 488 (Abcam, ab150077) in 1:1,000 dilution was used. The samples were visualised with FLUOVIEW FV10i confocal microscopy.

### Statistical analysis

Accurate and erroneous predictions were used to calculate the accuracy and error rate of classification of the cases to no regression, partial regression and total regression categories according to [14]. To provide further evidence that the results of the classification process is highly significant we calculated Matthews Correlation Coefficient (MCC) according to [15].

## Results

### Large RPB1 cytoplasmic foci are apparent in non-regressive invasive carcinoma of NST cells

We aimed to investigate the contribution of co-translational protein aggregation to the chemoresistance of invasive carcinoma of NST. Therefore, immunofluorescent microscopy was performed to follow the cytoplasmic aggregation of RPB1, the largest subunit of RNAPII in biopsy samples, taken from patients before the administration of neoadjuvant chemotherapy (Figure 1, S1). Compared with patients who responded to the therapy with either partial or complete tumor regression, those with tumors showing no regression in tumor size following chemotherapy had numerous cytoplasmic RPB1 aggregated foci in their first biopsy samples.

**Figure 1.**
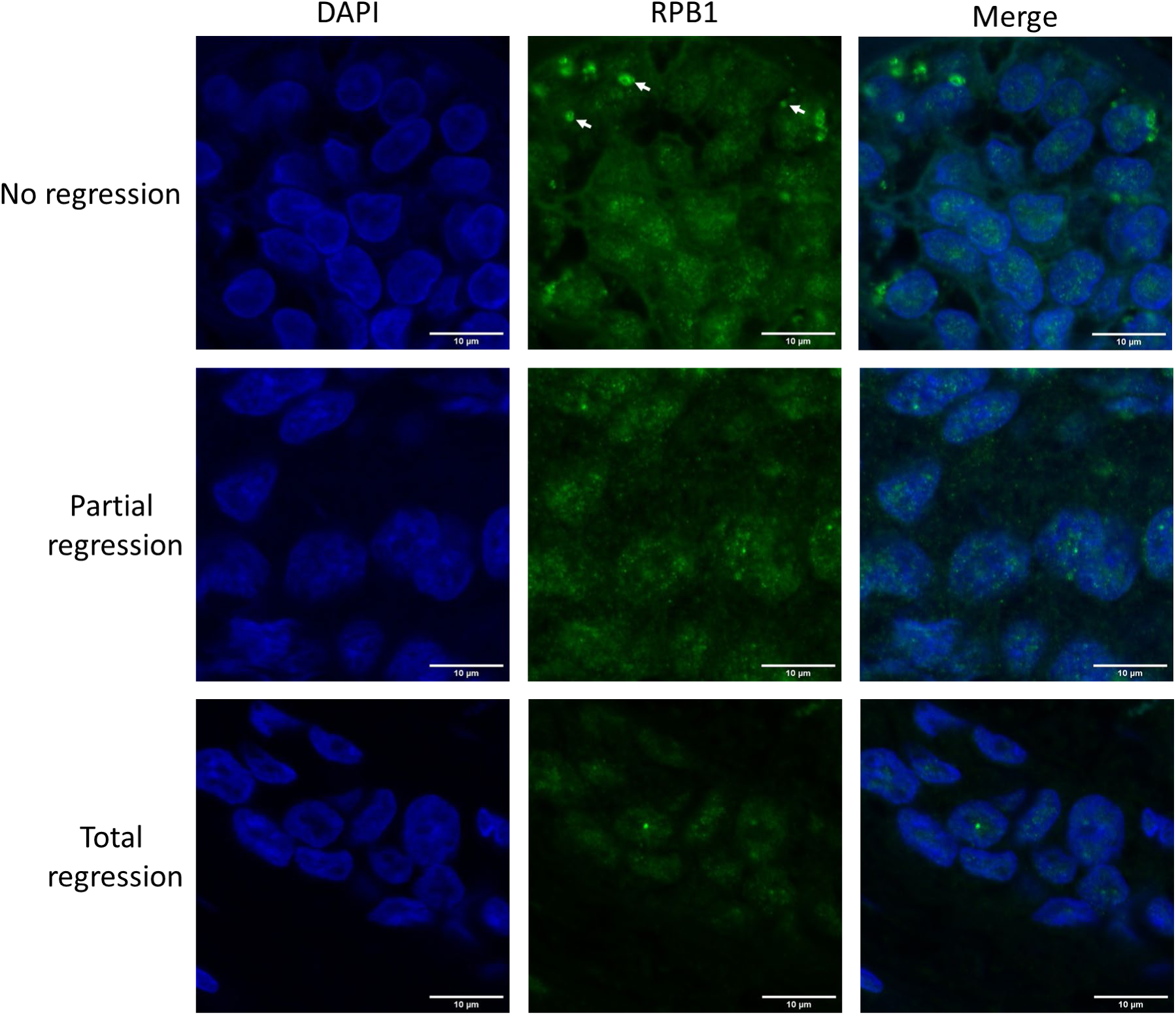
RPB1 appears in cytoplasmic foci in invasive carcinoma of no special type (NST) cells resistant to neoadjuvant chemotherapy. Immunofluorescent images of one representative case are shown from each of the three phenotypic categories as indicated on the left. In the cytoplasm of chemo resistant tumor cells — showing no regression after therapy — RPB1 foci are apparent (three examples are shown with white arrows). Nuclei are revealed using DAPI in cyan. RPB1 is shown in green, and the merged images of the two staining are highlighted. Scale bar: 10 μm.

As a control to test our sample preparation and staining protocol, we followed a modified nuclear RPB1. The fifth serine at the C-terminal heptapeptide repeated domain of RPB1 becomes phosphorylated at promoters by the basal transcription factor TFIIH [16]. We could detect only nuclear RPB1 as expected, with the phospho-5-serine modified RPB1 recognizing antibody in all three phenotypic categories (Fig. S2), clarifying that the cytoplasmic staining is not an artifact of sample preparation, fixation, or staining.

After investigating the samples with known regression phenotypes, we tested the use of the RPB1 cytoplasmic phenotype screen in predicting the outcome of therapy by categorizing samples into the three possible outcome groups in a blind experiment. Of the 13 investigated samples, we have managed to correctly categorize 10 (Fig. S1). Prediction accuracy of no regression, partial regression and total regression cases were above 84%, 76% and 92% respectively, as calculated according to [14] (Table S1). Matthews correlation coefficient (MCC) describes the relationship between the classification model, a perfect classification (which has a MCC of 1), random guessing (Which has a MCC of 0), and a total disagreement, meaning there are only false positives and negatives (which has a MCC of - 1)[15]. MCC of the prediction of no regression, partial regression and total regression cases were 0.675, 0.5 and 0.8216 respectively giving credit to the classification method based on following Rpb1 cytoplasmic aggregation phenotypes (Table S1).

### Cytoplasmic RPB1 foci are apparent sporadically in renal cell carcinoma cells, although not in the cells of surrounding tissues

We investigated another type of cancer for a similar phenotype to determine if the presence of cytoplasmic RPB1 foci is a specific phenotype observable only in invasive carcinoma of NST. We chose to study clear cell renal cell carcinoma samples from patients who underwent radical or partial nephrectomy on samples analyzed previously using fluorescent microscopy (Figure 2, Table S2) [13].

**Figure 2.**
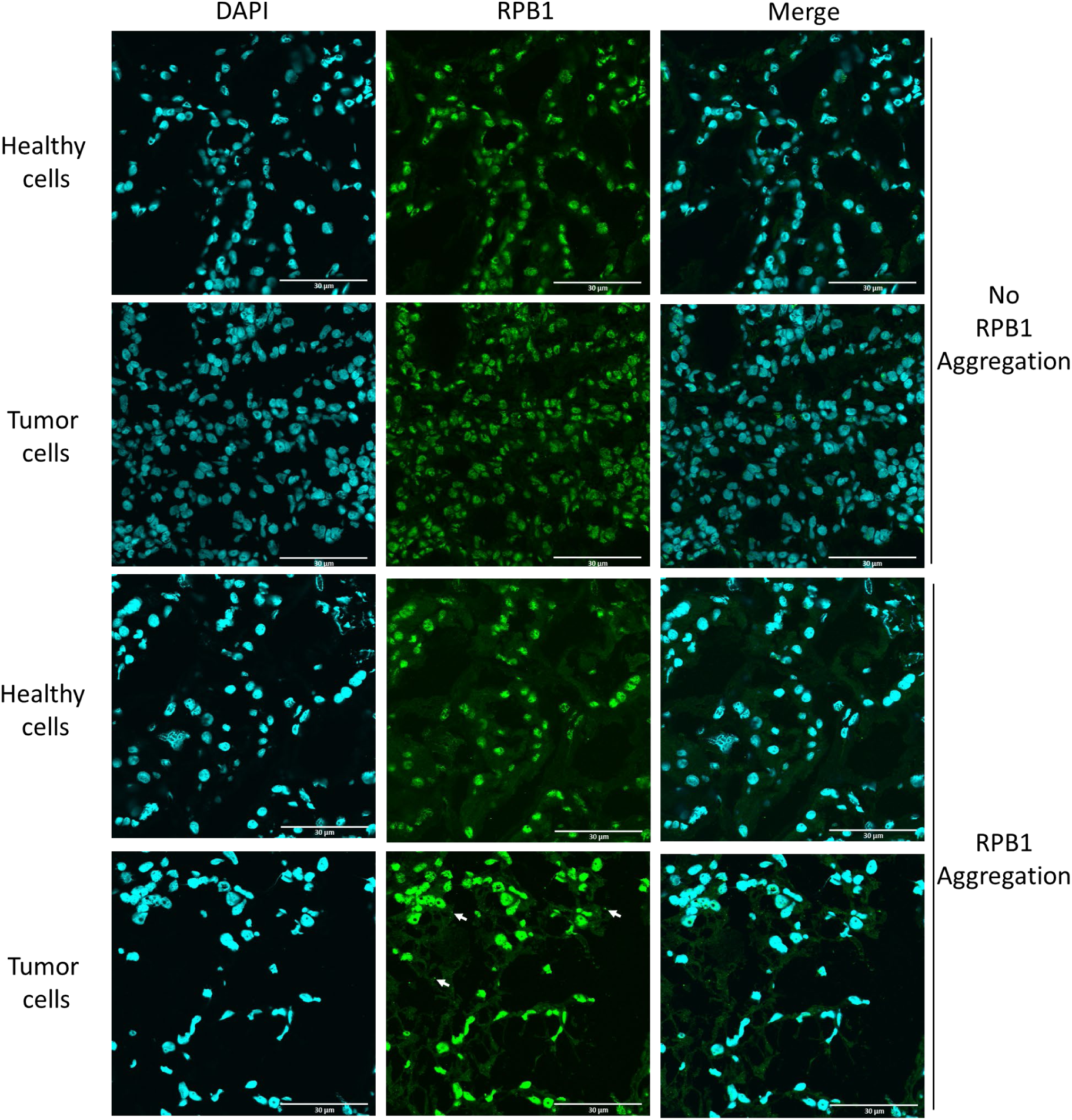
RPB1 aggregates sporadically in clear cell renal cell carcinoma. Fluorescent images of one representative case of clear cell renal cell carcinoma are shown, wherein RPB1 shows no cytoplasmic aggregation, and RPB1 foci are apparent (three examples are shown with white arrows). We have identified 18 patients where tumor-specific aggregation of RPB1 is detectable in surgically removed renal samples in a cohort of 30 patients (Table S2). Nuclei are revealed using DAPI shown in cyan. RPB1 is shown in green, and the merged image of the two staining are highlighted. Scale bar: 30 μm.

Here, we investigated both the malignant cells and the surgically removed surrounding healthy tissue. According to our results, RPB1 cytoplasmic foci were observable in some clear cell renal carcinoma cells, although not in surrounding healthy renal tissue. Thus, we clarified that RPB1 cytoplasmic accumulation is not specifically observed in invasive carcinoma of NST; however, based on the analysis of 30 patients and 100 cells, it appears to be tumor-specific in renal tissues. Cytoplasmic RPB1 foci were discovered in the tumor samples of 18 patients in the cohort.

## Discussion

Reoccurring failure in protein synthesis can lead to the loss of function of specific tumor-suppressor proteins. It is easy to see that a mutation affecting a ribosomal protein (RP) that is situated exactly where the correct interaction with the tRNA, for example, the decoding of the mRNA, normally takes place can lead to such failure and lead to the formation of aberrant or non-functional proteins. RPL5 and RPL10 are located in sites where decoding takes place inside the ribosome and exhibit mutations with considerable frequency in tumor cells [17]. A recent study has reported that circulating tumor cells isolated from patients with breast cancer exhibited increased ribosome biogenesis (RiBi) and RP translation, pointing toward an altered function of their translation machinery [18]. NOT1 of the CCR4–NOT complex is associated with RiBi and RP mRNAs facilitating their translation in yeast [19, 20]. Furthermore, NOT1 has shown to become engaged with stress granules and is important in the formation of assemblysome and mRNA condensates [4,12]. Cytoplasmic mRNA-containing condensates represent the reservoirs of transcription independently expressible gene products. One possible explanation of how tumor cells become resistant to transcription blockade by epirubicin is that they contain stored mRNA in condensates acting as a reservoir to maintain gene expression; simultaneously, mRNA decay initiated by the major deadenylase CCR4 function is altered in them. Consequently, the loss of transcription is buffered by other gene expression processes. Even the loss of protein degradation may be beneficial if the transcription is limiting. Remarkably, the CCR4–NOT complex contains the major deadenylase of eukaryotic cells and, as a chaperone platform, it has been revealed to be involved in the assembly of RNAPII and proteasome [4,9]. Although it remains elusive, it is tempting to assume that frequent aggregation of RPB1 in tumor cells resistant to the transcription inhibitor epirubicin may be a consequence of failure in the CCR4–NOT dependent co-translational folding of RPB1. Notably, other functions of the CCR4–NOT complex that may become more pronounced in tumor cells at the expense of its role in RNAPII assembly are exactly roles that have the potential to supplement the loss of transcription, including its role in mRNA condensate formation and RP and RiBi translation [12,19]. Increased RiBi provides the basic machinery for translation, which has the potential to buffer the loss of the other major gene expression process under epirubicin-induced transcription stress.

Considering its predictive significance, we find it urgent to report that based on our results, and the blind test predicting no regression cases above 84% accuracy, RPB1 has a tendency to aggregate in epirubicin-resistant tumor cells. The phenotype is easy to follow from biopsy samples immediately after cancer diagnosis. We propose that patients with cytoplasmic RPB1 foci in their biopsy samples should consider the high risk of ineffective, but toxic treatment with transcription blockers along with the time loss and choose surgery instead of neoadjuvant chemotherapy.

Moreover, we report that RPB1 aggregation is not invasive carcinoma NST specific; however, it may be ubiquitous as we detected this phenotype in 18 patients in a cohort of 30 diagnosed with clear cell renal cell carcinoma. The fact that the phenotype was not apparent in the healthy surrounding tissue in any of the surgically removed renal samples indicates that RPB1 cytoplasmic aggregation might be a consequence of an oncogenic mutation rather than present before malignant transformation in healthy renal tissue of susceptible patients.

## List of abbreviations

P-body: Processing Body
RPB1: RNA Polymerase B (II) subunit 1.
HSP90: Heat Shock Protein 90
NOT1: Negative on TATA-less 1
CCR4: Carbon Catabolit Repression 4
NST: No Special Type
IPA: Isopropanol
BSA: Bovine Serum Albumin
GAM: Goat Anti Mouse
GAR: Goat Anti Rabbit
PBS(T): Phosphate Buffered Saline with Tween 20
ACC: Accuracy
ERR: Error Rate
MCC: Matthews Correlation Coefficient
RPL5: Ribosomal Protein L5 (L:large subunit)
RPL10: Ribosomal Protein L10 (L:large subunit)
RP: Ribosomal Protein
RiBi: Ribosome Biogenesis

---

**Supplementary figure 1.**

One representative fluorescent image from each of the studied invasive carcinoma of NST biopsies is listed and categorized according to response to neoadjuvant treatment. “No regression”, “partial regression” and “total regression” cases are inside red, yellow and green boxes, respectively. Case numbers for the internal record are shown in the middle. Eleven cases with known phenotypes are on the left, and additional 13 cases who participated in the blind test are on the right. Case numbers with different colors as the bracket indicate the cases that have been categorized falsely (three cases), and the color of the number represents the falsely assumed category, respectively. Arrows indicate three examples for RPB1 foci in each of the images taken from tumors belonging to the real (left) or assumed (right) “no regression” phenotypic category. Scale bars for images on the left: 10 μm and right: 30 μm.

**Supplementary figure 2.**

Antibodies recognizing nucleus engaged RPB1 phosphorylated at the fifth serine of its heptapeptide C-terminal repeats (P5-RPB1-CTD) reveals cytoplasmic staining in none of the three studied phenotypic categories as indicated on the left. Nuclei are revealed using DAPI shown in cyan. RPB1 is shown in red, and the merged image of the two staining is highlighted. Scale bar: 10 μm.

**Supplementary table 1.**

Confusion matrixes of the blind test. Accurate and erroneous predictions are highlighted and are used to calculate the accuracy (ACC) and error rate (ERR) of classification of the cases to no regression, partial regression and total regression categories according to [14]. Matthews Correlation Coefficient (MCC) is highlighted as calculated according to [15]. The complete confusion matrix and the confusion matrixes for each regression categories are highlighted on separate pages.

**Supplementary table 2.**

Clear cell renal cell carcinoma sample numbers are representing tumors where RPB1 has appeared or not in cytoplasmic foci respectively. The sample numbers are as described in

[13].

